# Interaction paths promote module integration and network-level robustness of spliceosome to cascading effects

**DOI:** 10.1101/302570

**Authors:** Paulo R. Guimarães, Mathias M. Pires, Maurício Cantor, Patricia P. Coltri

## Abstract

The functionality of distinct types of protein networks depends on the patterns of protein-protein interactions. A problem to solve is understanding the fragility of protein networks to predict system malfunctioning due to mutations and other errors. Spectral graph theory provides tools to understand the structural and dynamical properties of a system based on the mathematical properties of matrices associated with the networks. We combined two of such tools to explore the fragility to cascading effects of the network describing protein interactions within a key macromolecular complex, the spliceosome. Using *S. cerevisiae* as a model system we show that the spliceosome network has more indirect paths connecting proteins than random networks. Such multiplicity of paths may promote routes to cascading effects to propagate across the network. However, the modular network structure concentrates paths within modules, thus constraining the propagation of such cascading effects, as indicated by analytical results from the spectral graph theory and by numerical simulations of a minimal mathematical model parameterized with the spliceosome network. We hypothesize that the concentration of paths within modules favors robustness of the spliceosome against failure, but may lead to a higher vulnerability of functional subunits which may affect the temporal assembly of the spliceosome. Our results illustrate the utility of spectral graph theory for identifying fragile spots in biological systems and predicting their implications.

## 1. Introduction

Multiple biological systems are characterized by networks^1,2^. Genes form regulatory networks^3,4^, proteins are connected through a network of paths^5,6^, individuals are embedded in social networks^7^, and species are linked to each other in food webs^8^. In the past two decades, we have learned about the main structural aspects of multiple biological networks^8-11^. Simultaneously, a wide range of empirical and theoretical studies explored the dynamical implications of network structure^5,12-14^.

Within the cell, the structure of protein networks may provide information about the underlying processes shaping the organization and function of macromolecular complexes and may affect the vulnerability of these macromolecular processes to distinct types of perturbations^15-18^. Mutations may lead to non-functional proteins that in some cases will imperil primal functions, leading to the death of the cells or affecting tissue functioning, causing multiple types of diseases^19,20^. Alternatively, some mutations may not impair spliceosome function but, on the contrary, stimulate one specific step of splicing. For instance, despite the importance of PRP8, a protein involved in the formation of the spliceosome catalytic core, some mutations on yeast PRP8 affect splicing efficiency and fidelity, but do not impair spliceosome formation^21,22^. These examples might indicate that the underlying protein network is robust. Hence, a key challenge to the study of protein networks is understanding if and how network structure affects the fragility of protein interactions to perturbations.

One of the main venues to explore the role of network structure to system functioning is via mathematical modelling. One challenge of mathematical modelling of complex systems is the parameterization. A way to circumvent this challenge lies in the fact that multiple dynamical processes in complex systems are shaped by the architecture of the underlying network^12^. By focusing on the possible role of network structure on dynamics, one can assess the consequences of particular interaction patterns in spreading perturbations through the system. To this end, spectral graph theory is a powerful tool^23-26^. Spectral graph theory is the study of the eigenvalues and eigenvectors of matrices that describe or are associated with the networks. Specifically, the distribution of eigenvalues (spectra) of the adjacency matrix and the Laplacian matrix, two types of matrices associated with a given the network, contain information on the potential role of network structure in favoring or constraining cascading effects in distinct systems^23-25,27,28^.

Here, we explore the fragility of the network describing protein interactions within a key macromolecular complex, the *Saccharomyces cerevisiae* spliceosome (Fig. 1). Spliceosome catalyzes splicing, an essential process for gene expression regulation in eukaryotic cells^29^. Using tools from spectral analysis, we investigated the vulnerability of the spliceosome network to cascading effects. Since paths linking proteins indirectly provide propagation routes for cascading effects, we first used results related to the spectra of the adjacency matrix to estimate how these indirect paths are distributed within the spliceosome network. Then, we used the spectra of the Laplacian matrix associated with the spliceosome network to estimate the vulnerability of the network to cascading effects, in which changes in the state of a protein (mutations, unfolding, misprocessing, malfunctioning) may propagate across the network. Finally, we simulated cascading effects in these networks using a simple mathematical model and compared the simulated dynamics with the analytical predictions derived using spectral graph theory.

**Fig 1.**
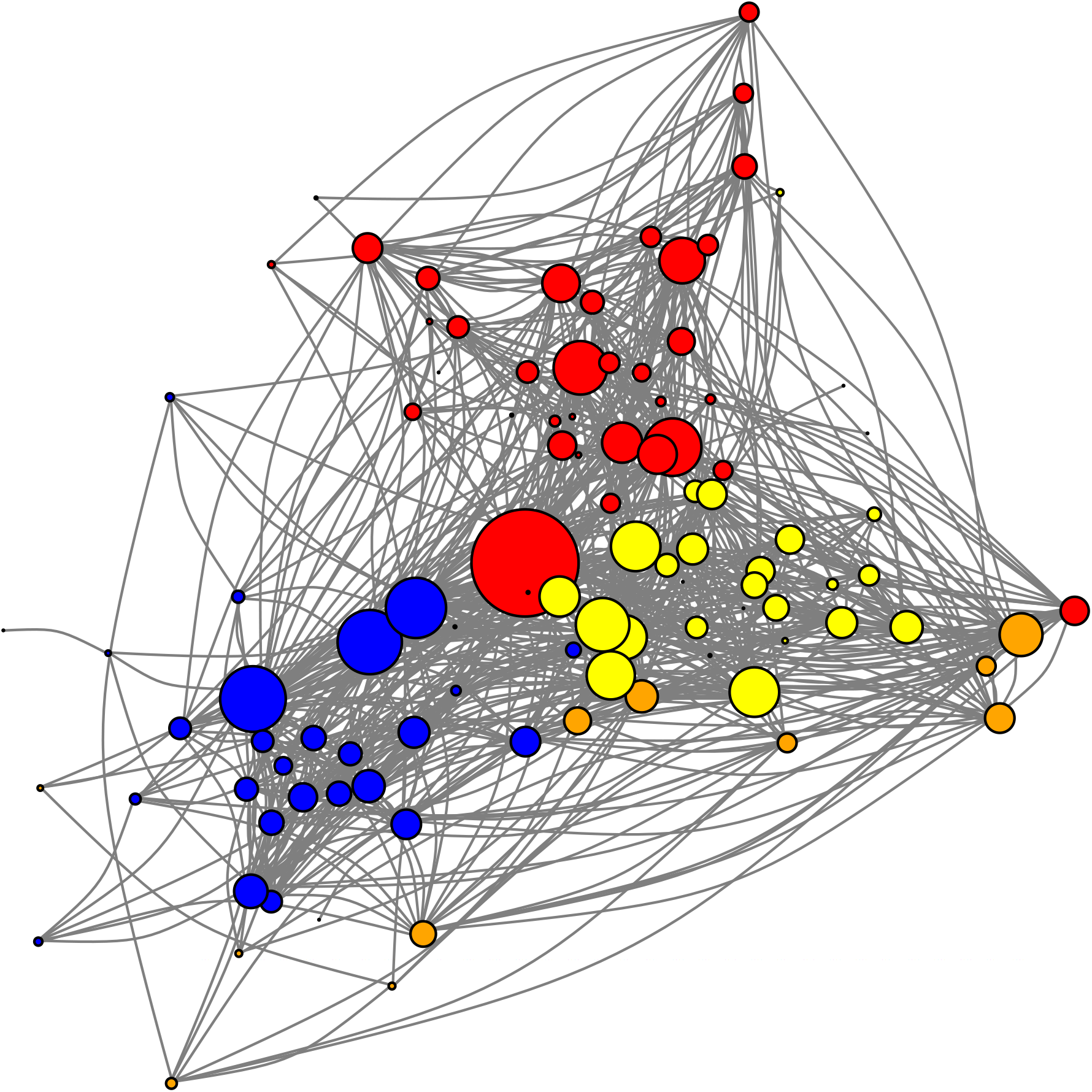
A network describing protein-protein interactions (links) distributed among 103 proteins (nodes) from the spliceosome of *Saccharomyces cerevisiae*. Links are defined based on the reliability of the evidence favoring protein-protein interaction (see text for further details). Node size is proportional to the number of interactions per protein and proteins with the same color are components of the same module in the network^31^. Modules represent groups of proteins that are more densely connected to each other than to other proteins in the network.

## 2. The spliceosome network

The spliceosome is composed of 5 snRNAs (small nuclear RNAs U1, U2, U4, U5 and U6) and more than 100 proteins. It is assembled on every intron during transcription after identification of important conserved sequences on the pre-mRNA. As a consequence, proteins and snRNAs interact sequentially and this ordered assembly is important to create a catalytic center responsible for the splicing reaction. The structure of this macromolecular complex suggests proteins and snRNAs are organized in different sub-complexes or modules, important for formation of the complex catalytic core^30^. These modules are sequentially rearranged as the spliceosome assembles, promoting protein-RNA interactions that will lead to activation of this complex. Therefore interaction between these modules is important to create the an active catalytic center. We analyzed the protein network of the *S. cerevisiae* spliceosome, formed by 103 proteins and their pairwise interactions^2,31^. Proteins and putative protein-protein interactions were recovered from the STRING database^32^. Pairwise interactions can be inferred using different approaches and these approaches may provide different levels of support to a putative pairwise interaction between two proteins^2,31^. A reliability score was assigned (varying from zero to one) to each putative protein-protein interaction according to the level of evidence suggesting the interaction to occur and provided by different experimental approaches^33^. The proteins are depicted as nodes and we assume there is a link connecting two proteins if the level of support to that putative interaction is higher than 0.50 (additional details at ^2,31^). This level of support represents a heuristic cutoff since lower cutoffs imply, by definition, in the record of weakly supported protein-protein interactions and higher cutoffs do not imply in structural changes to network structure^31^. Nevertheless, we performed a sensitivity analysis using a lower cut-off (0.15), since previous work showed that these two cutoffs (0.15 and 0.50) represent the two qualitative distinct network structures for the spliceosome^31^. The results assuming the lower cutoff were not qualitatively distinct from the patterns recorded for the higher cutoff (see Supporting Information) and we used the higher cutoff for the next analysis. The dataset is available at ^2^ and all the analyses were ran using MATLAB scripts that are available upon request.

We recorded 881 interactions, a fraction of all possible interactions actually recorded (the connectance) of *C* = 0.168. In this network, each protein has, on average, 17.11 ± 13.04 interactions^31^. In this sense, the spliceosome network is similar to other protein networks in which most proteins interact with a few other proteins and there is a small set of highly-connected proteins^15,34,35^. Proteins with higher molecular weights, as PRP8 and BRR2, had a higher number of interactions. Consistently, proteins with lower molecular weights, as CWC24 and CWC15, showed fewer interactions^31^. Previous studies showed that this spliceosome network is nonrandomly structured, combining patterns of nestedness and modularity. Nestedness occurs when proteins with fewer interactions and highly connected proteins interact, and at the same time, the highly connected proteins interact with each other^2^. Modularity is defined by the observation of cohesive groups of proteins that interact more with each other than with the rest^31^ (See also Table S1). Accordingly, the spliceosome network shows high levels of clustering, as indicated by a cluster coefficient much higher than the observed connectance (Table S1). The modular structure of the spliceosome network analyzed here was previously characterized in four modules identified using the maximization of the *Q* index of modularity in a simulated annealing framework^31^ (Fig. 1). These modules are associated with functional subunits of the complex (additional details in ^31^). Such structural patterns are robust when analysing spliceosome networks using a distinct dataset for *S. cerevisiae* spliceosome and are similar to the patterns recorded in human spliceosome network^31^.

## 3. Computing indirect paths among proteins

One crucial implication of the network structure of biological networks is the emergence of paths connecting distinct elements of the network directly or indirectly^27,36^. In the spliceosome network, these paths connect otherwise spatially and temporally isolated pairs of proteins and, on average, pairs of proteins are at only two degrees of separation to each other (estimated by the average shortest path length, Table S1). Given that cascading effects may propagate through such paths^36^, quantifying how distinct proteins are connected in the network may inform on the fragility of the spliceosome to cascading effects. A path is defined as a set of nodes (here, proteins) and the links connecting them (here, protein-protein interactions), starting and finishing with nodes (Fig. 2A). The path length, *ζ*, is the number of links in a path. Note that we are using the broader definition of path (i.e., a walk), in which the same node or link may be represented multiple times in a path and, therefore, longer paths may be formed by a combination of smaller paths (Fig. 2A). As a consequence, the number of paths increases exponentially with path length, *ζ*, a phenomenon called path proliferation^36^.

**Fig 2.**
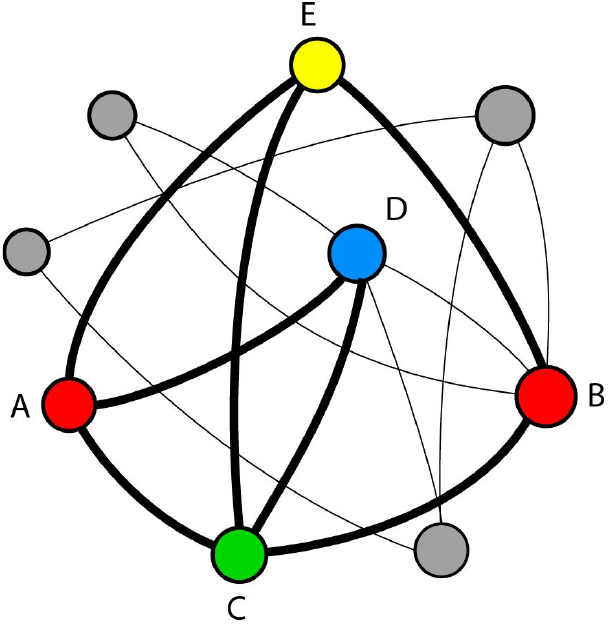
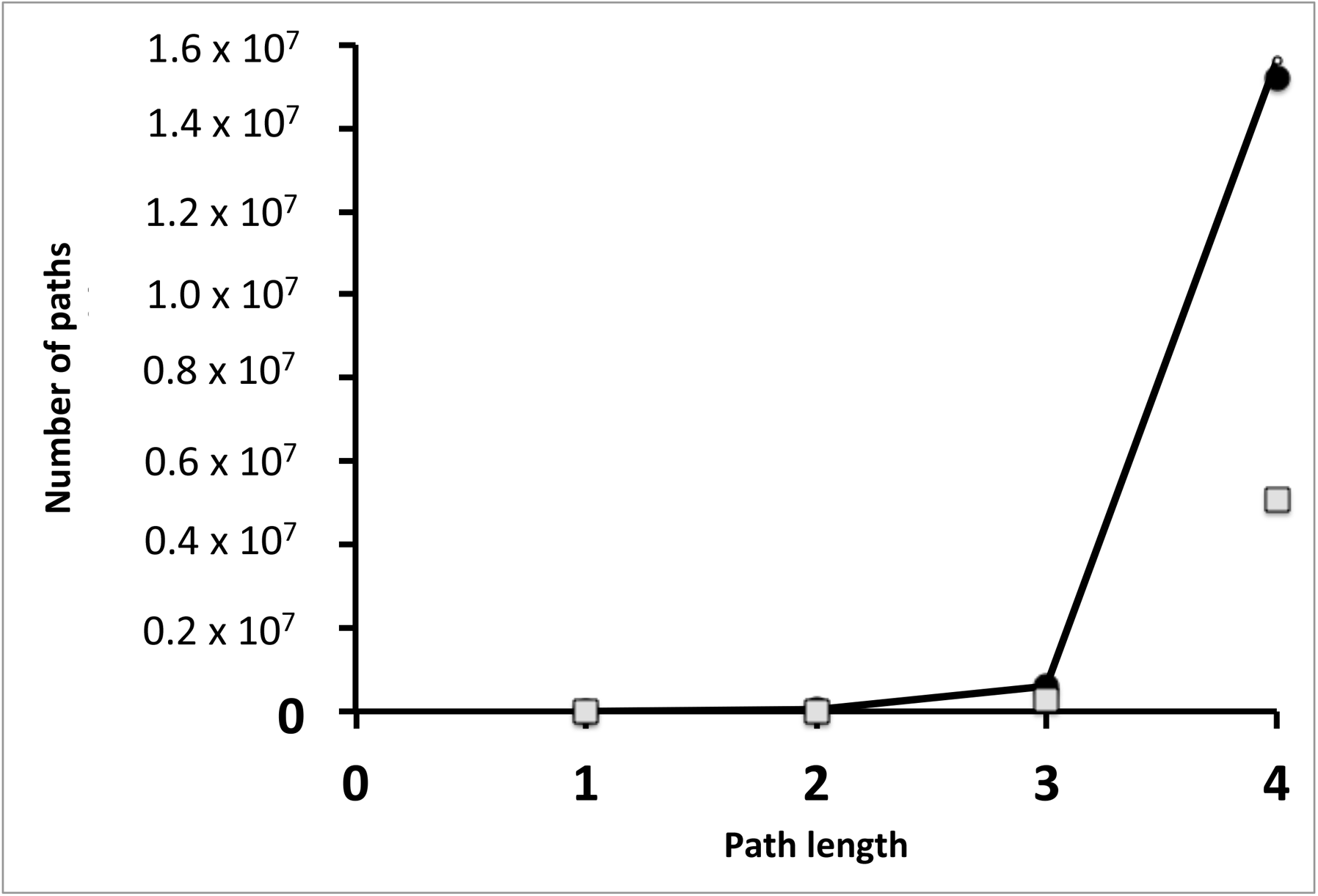
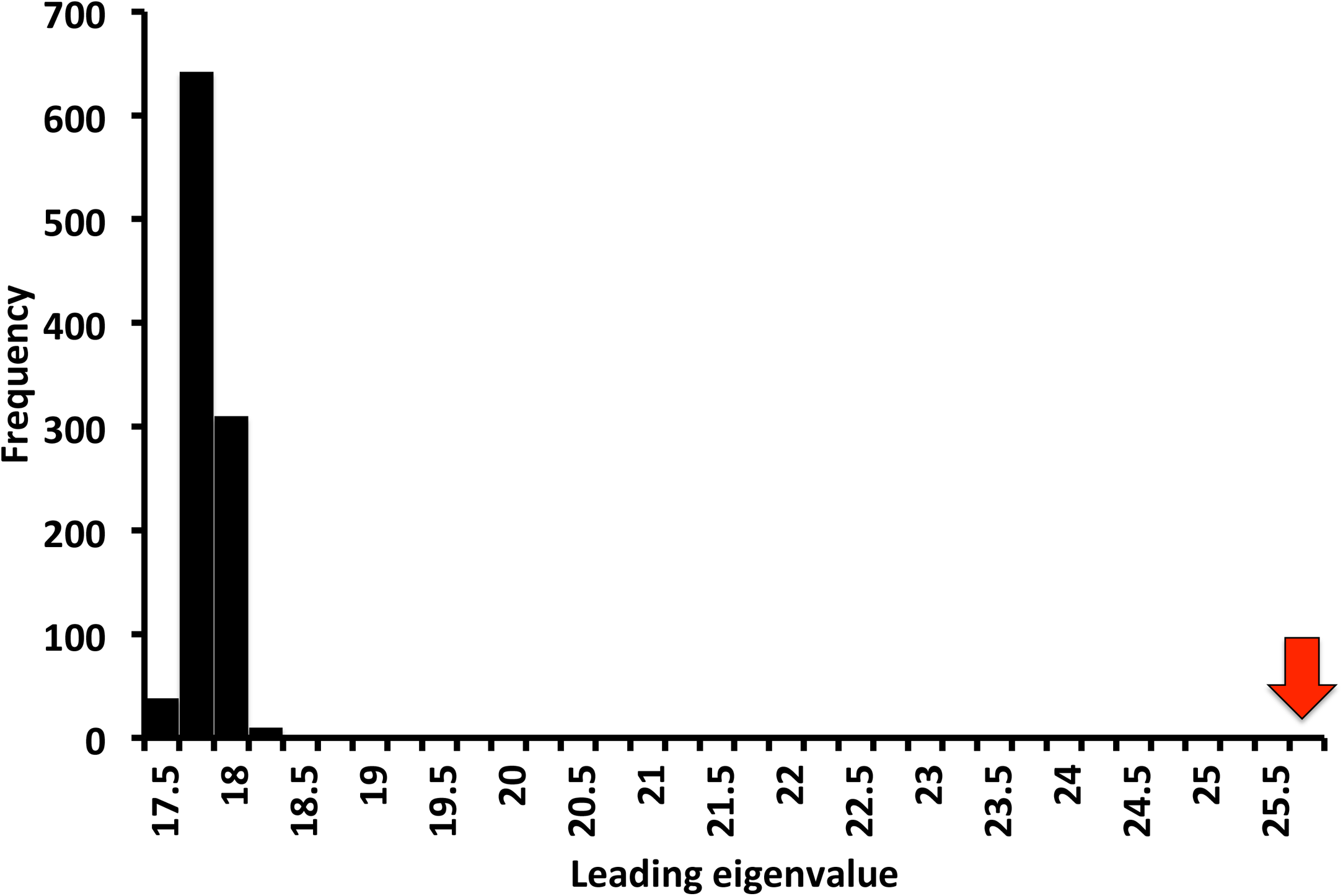
Paths in a network. A) A hypothetical network in which two proteins (A,B) are connected by multiple paths of different lengths, including A-C-B, A-E-B, A-D-C-B, A-E-C-B. B) The number of paths increases faster with path length for the spliceosome network (closed circles) than for random networks with the same number of proteins and protein-protein interactions (gray squares). The trendline describes the analytical prediction for the spliceosome network based on the leading eigenvalue of the adjacency matrix (see text for further details). C) The higher path proliferation of the spliceosome network is a consequence of the larger leading eigenvalue of its adjacency matrix (red arrow) when compared with the expected leading eigenvalue of random networks (black bars, 1000 random networks).

The higher the density of paths in a network with a given length *ζ* the higher the number of possible routes that allow perturbations to cascade through proteins in the network. Therefore, the rate of path proliferation is a useful statistic to describe the multiplicity of routes allowing cascading effects in a given network^36^. The rate of path proliferation is related to the eigenvalues of the adjacency matrix **A**^36,37^. In the adjacency matrix **A**, a given element *a_ij_* describes if the interaction between two proteins *i* and *j* occurs (*a_ij_* = *1*) or not (*a_ij_* = *0*). Although the rate of path proliferation is related to all eigenvalues, for large values of *ζ* the rate of path proliferation is governed by the leading eigenvalue of **A**, *λ_A_*, and the number of paths with a given length *ζ*, *ψ*^(*ζ*)^, is

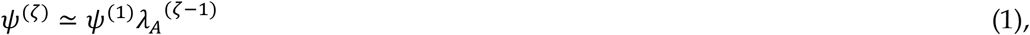

in which *ψ*^(*1*)^ is the number of protein interactions recorded in the spliceosome network. The higher the *λ_A_*, the higher the number of paths of a given length *ζ* connecting proteins in the network.

We first computed the leading eigenvalue of the spliceosome network, which is *λ_A_* = *25.84*. We used the leading eigenvalue to predict the accumulation of paths with the increase of *ζ* and we compared to actual accumulation of paths in the spliceosome network. The total number of direct interactions between pairs of proteins (paths of length *ζ* = *1*) in a network with *N* proteins is equal to 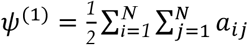. Likewise, the number of paths of length *ζ* = *2* can be estimated by computing **A**^*2*^. The element 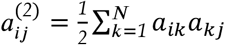, which is nonzero if there is at least one protein *k* interacting with both proteins *i* and *j* and 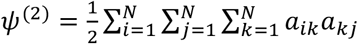. Hence, the number of paths of length *ζ* = *h* is the sum of all elements of matrix 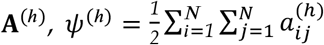. We computed the rate of path proliferation in the spliceosome network. The rate of path proliferation follows the analytical prediction derived from spectral graph theory even for short paths, *ζ* < *4* (Fig. 2B), although equation (1) represents a prediction for the number of paths assuming large *ζ*.

We now turn our attention to the role of the nonrandom structure of the spliceosome network in shaping the proliferation of paths. We computed the leading eigenvalues of 1,000 random networks with the same number of proteins and the same number of interactions but in which the probability that a protein *i* interacts with protein *j* is constant and equal to the connectance. Spectral graph theory predicts that for a random graph (an Erdos-Renyi graph) in which *NC* >> log (*N*) the expected value for the leading eigenvalue for a random network is λ_*R*_ = [*1* + *o(1)*]*NC*, in which *o*(*1*) is a function that converges to a value close to zero^38^. Thus, the expected leading eigenvalue for a random network with the same number of proteins and interactions is [*1 + o(1)*]*NC* ≃ *NC* = *17.274*. This prediction is close to the mean leading eigenvalue for our simulated random networks (17.94±0.12). Both the analytical prediction for random networks and the estimated value for simulated random networks are much smaller than the leading eigenvalue of the spliceosome network (*λ_A_* = *25.84*; Fig. 2C, *P* < 0.001). This difference in the leading eigenvalues of spliceosome network and random networks implies that, for a given path length, the empirical spliceosome network is much more connected than expected by chance. For example, for *ζ* = *3*, there are 602,250 paths connecting the spliceosome proteins in the network. In contrast, for a random network with a similar number of proteins and a similar number of protein interactions the expected number of paths of length *ζ* = 3 is just one third of the number of paths observed in the empirical network (282,810±3,644.1; ***P** < 0.001*). These results are consistent when we assumed a more conservative null model that maintain the heterogeneity in the number of interactions per protein (Supplementary Information).

## 4. Paths and modular structure

The spliceosome network has a modular structure, in which interacting proteins are concentrated into cohesive subgroups^31^. Such modular structure may counterbalance the role of paths in facilitating cascading effects. Since there are no disconnected components in the spliceosome network, there are paths connecting all pairs of proteins. However, modularity may prevent the homogenizing effects of indirect paths since most of the paths start and end in proteins of the same module^27,36^. If this is true, then cascading effects are expected to propagate among modules less efficiently and the consequences of indirect paths are expected to be greater locally, i.e., within modules, than between modules^36^.

The spliceosome network analyzed here has four modules^31^. We investigated if paths are more concentrated within modules than between modules. To compare the role of paths in connecting proteins from the same or from distinct modules, we computed the mean number of paths of a given length *ζ* starting and ending with proteins in the same module, 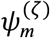, and starting and ending with proteins from distinct modules, 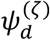. We then computed the ratio 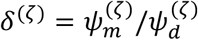. The direct interactions, *ζ* = *1*, are concentrated within modules: 36.03% of all possible interactions between proteins of the same module were observed, whereas just 8.94% of the possible interactions between proteins of different modules were observed, leading to *δ*^(*1*)^ = *4.03*. Because the modules are not totally disconnected, indirect paths connect proteins from different modules, leading to a reduction of *δ*^(*ζ*)^ as *ζ* increases (Figure 3). That said, the effects of the modular structure still affect the path distribution. For example, when *ζ* = *2*, there were twice as much paths per pairs of proteins within modules than between modules (*δ*^(*2*)^ = *1.99*). The values of *δ*^(*ζ*)^ converge to values close to one as *ζ* increases, but even for very large values of *ζ*, there is still a small concentration of paths within modules when compared with paths between modules (*δ*^(*ζ→∞*)^ = *1.02*). Thus, although the spliceosome network structure favors a higher density of paths than expected in random networks, the modular structure implies that most of these paths are concentrated within rather than between modules.

**Fig 3.**
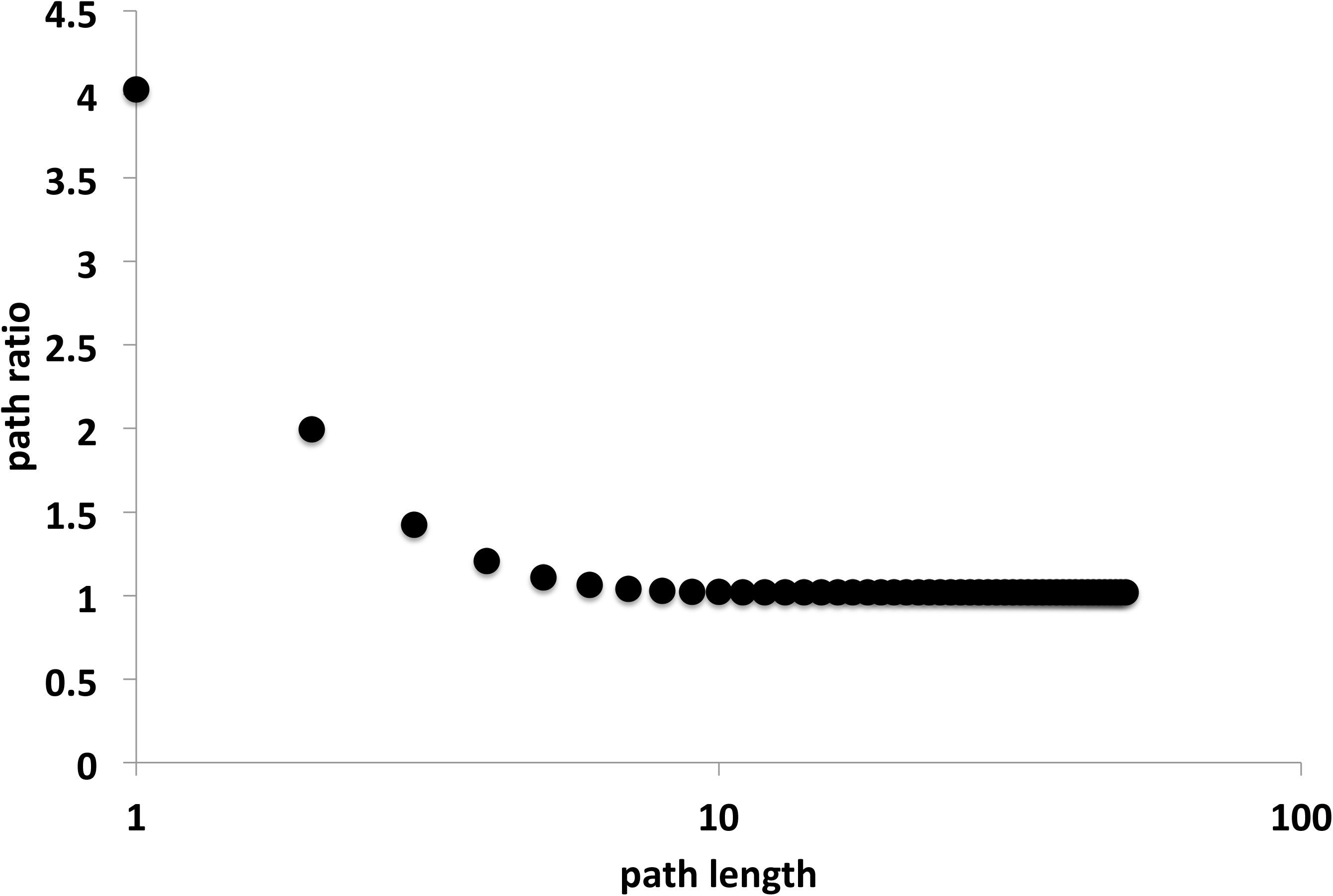
Paths within and across modules. Path ratio is the ratio between the number of paths starting and ending with proteins from the same module to the number of paths starting and ending with proteins from different modules. The concentration of paths within modules decays with path length. Nevertheless, even for long paths most of paths connect two proteins from the same module (path ratio >1).

## 5. How likely are cascading effects to spread across the network?

We showed that despite a certain level of inter-module connectedness, most of the paths are concentrated within the modules of the spliceosome network. Modularity has been shown to limit the cascading effects of perturbations^39,40^. Therefore, a fundamental problem is to describe how isolated the network modules are. There are multiple approaches to measure the isolation of modules. The spectral analyses of the Laplacian matrix is an approach directly rooted in the implications of connectivity to the dynamics within networks, e.g., network flow, homogenization of states, and synchronization^41^. The Laplacian matrix, *L*, is defined in such way that if *i=j*, 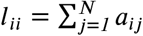, and if *i* ≠ *j*, *l_ij_* = −*aij*. Note that if *a_ij_* = 0 than *l_ij_* = 0. The spectral properties of the Laplacian matrix inform on the connectivity of groups within networks^24^. For example, the number of zero eigenvalues inform the number of isolated groups of proteins within a network (i.e., the network components). In a connected network, in which there are paths connecting any pairs of proteins, there is a single zero eigenvalue. In this case, the smallest non-zero eigenvalue, also called algebraic connectivity or Fiedler number, describes how well connected are modules within networks.

We computed the algebraic connectivity of the spliceosome network and compared it with that of 1,000 random networks with the same number of proteins and the same number of interactions. The algebraic connectivity of the spliceosome network is *λ*_*L*_ = *0.689*, a value much smaller than expected for a similar random network (7.15 ± *0.90, P* < *0.001*, Figure 4a). Therefore, the connectivity among modules in the empirical spliceosome network is much weaker than expected for a random network with the same number of interactions and proteins. These results hold for analysis assuming a more conservative null model (Supplementary Information).

**Fig 4.**
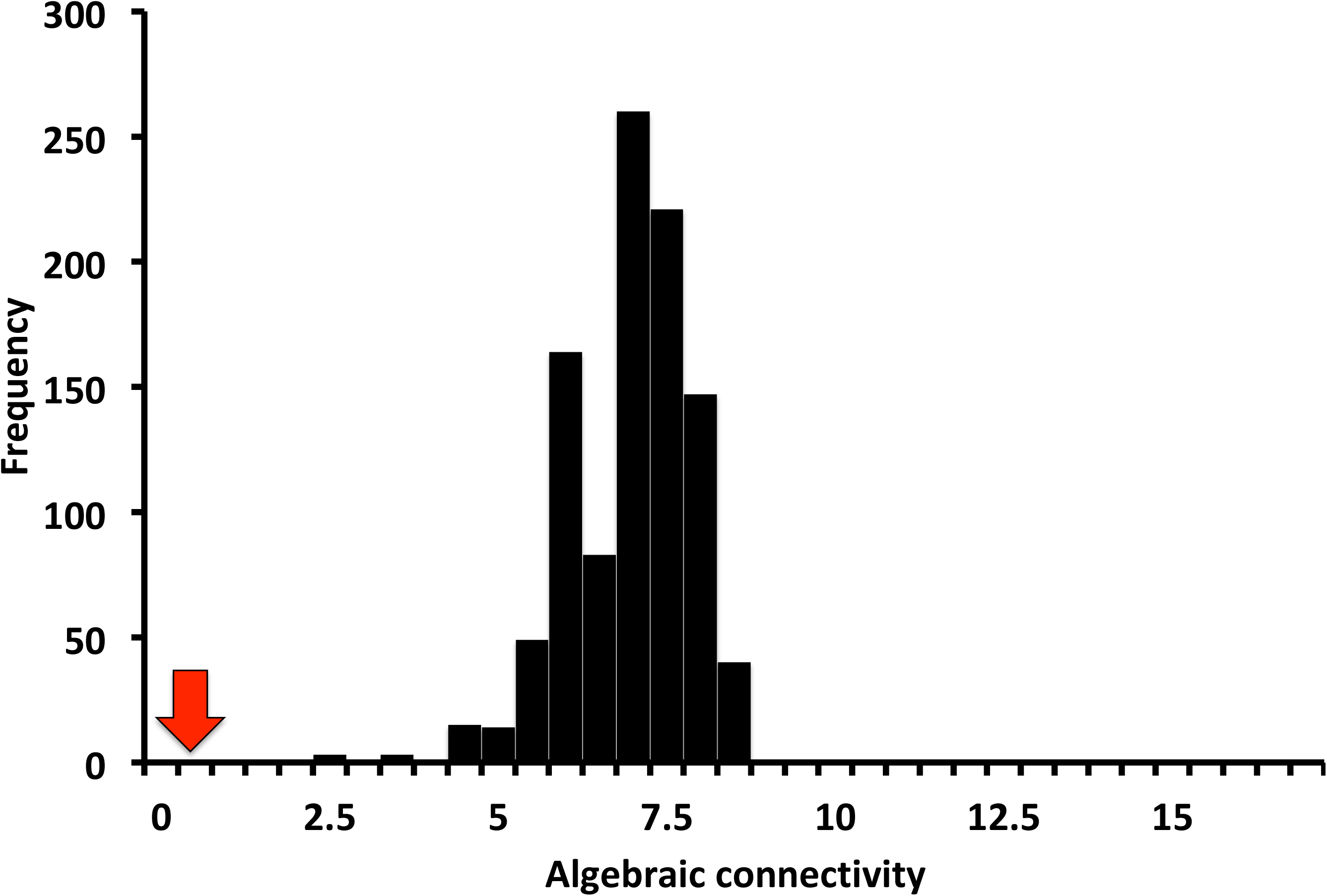
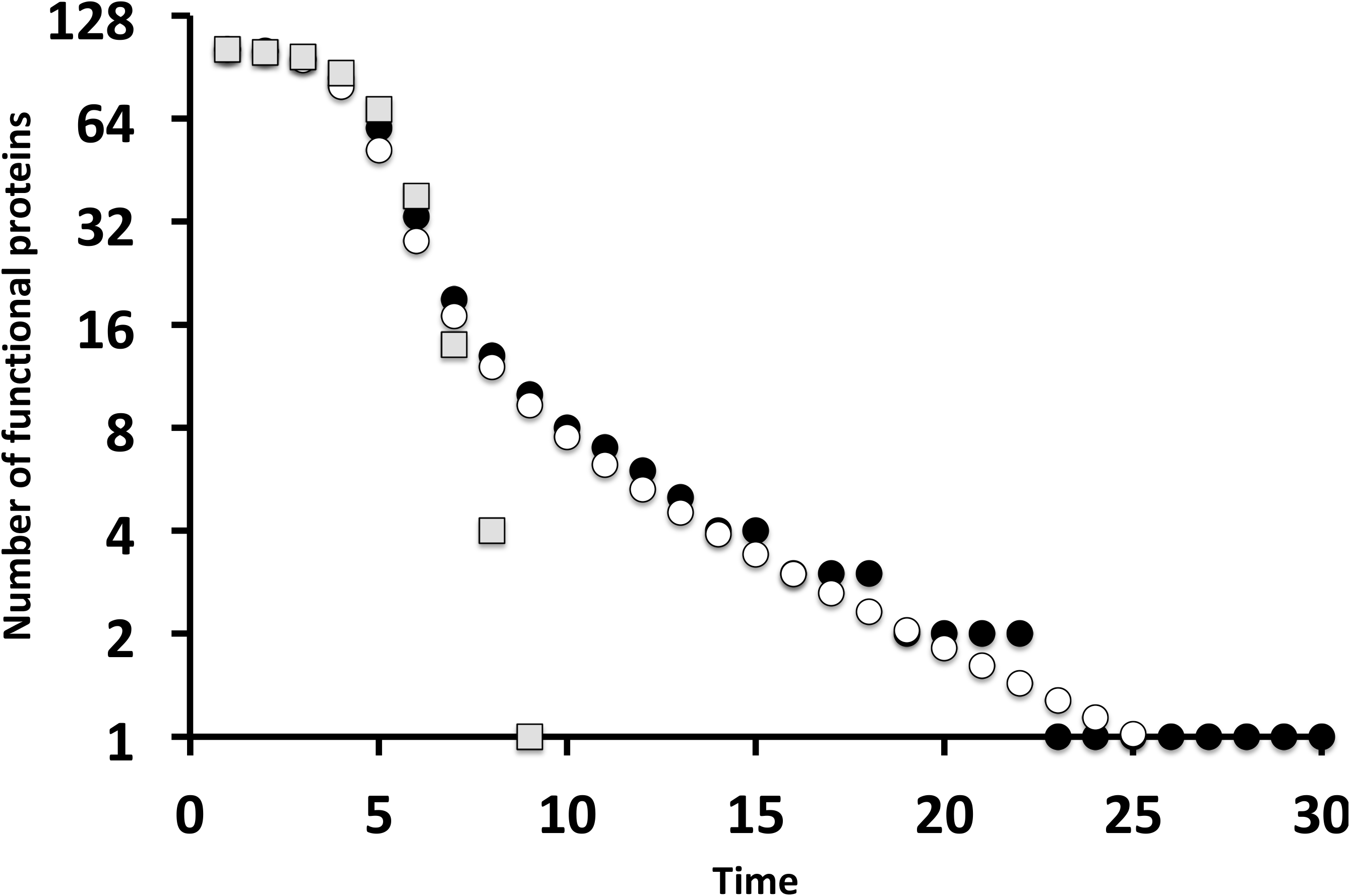
Cascading effects in the spliceosome network. A) Spectral graph theory predicts that the rate of spreading of cascading effects will be governed by the eigenvalues of the Laplacian matrix, especially the algebraic connectivity (see text for further details). The spliceosome network show lower algebraic connectivity (red arrow) when compared with the expected algebraic connectivity of random networks (black bars, 1000 random networks). B) The time-to-collapse of the spliceosome network in a minimal mathematical model of failure spreading. Each closed circle is the median number of functional proteins (1000 simulations) per time. Each open circle is the analytical prediction of the number of functional proteins per time derived from a mean-field approximation of the model (1000 simulations). Grey squares represent the median number of functional proteins in random networks (1000 simulations).

The spectral properties of the Laplacian matrix also affect how fast cascading effects impact the network by means of approaches derived from the study of flow of networks and of the emergence of synchronization. We used a set of difference equations to explore the consequences of the spectral properties of Laplacian matrix for the dynamics of the system. This minimal model does not aim to reproduce the dynamics of a given biological process in detail, but to explore the potential role of network structure in shaping cascading effects in very simple dynamics. Our minimal model assumes that there is a state associated to a given protein *i*, *ϕ*_*i*_, and the dynamics describe how fast direct and indirect effects lead to the homogenization of state values. The state *ϕ*_*i*_ may assume two different values, *ϕ*_*i*_ = *1* if the protein performs its function in the spliceosome and *ϕ*_*i*_ = *0* if the protein is mutated in a nonfunctional form or if the malfunctioning of interacting proteins lead to a cascading effect that inhibits the performance of an otherwise functional protein. We assume that the probability of a given protein to be affected by nonfunctional interacting proteins in a given time step is:

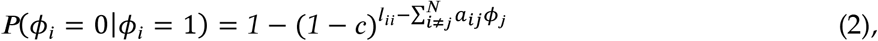

in which *c* is a constant between zero and one that controls the propagation of cascading effects. In this model, given enough time, all the proteins lose their functionality, so the spliceosome becomes dysfunctional. In reality, not all errors and mutations may lead to the collapse of the spliceosome^20,42^. However, by estimating how fast the functionality of proteins collapse due to errors, this minimal model allows estimating the effect of network structure in favoring or preventing cascading effects, without the complexity of real, empirical dynamics of biological processes. The faster the convergence in state values, the higher is the contribution of network structure to fuel cascading effects. The dynamics of the model can be approximated by a mean-field, continuous model:

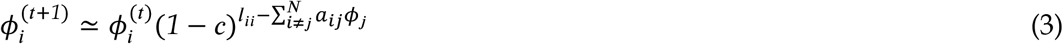

In which 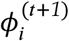 is a continuous variable. Under the assumption that most proteins are still functional, we can approximate (3) to:

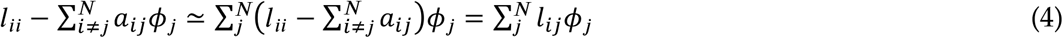

Substituting (4) in (3) the equation (3) can be generalized to all proteins of the network. In matrix form, the resulting set of equations represent all the difference equations of the network is:

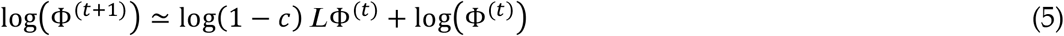

in which Φ^(*t*)^ is a vector with the states of all proteins at time *t*. Note that *c* could assume distinct values for each protein but for the sake of simplicity we used the same value for all proteins without any loss of generality. Therefore, the Laplacian matrix informs how fast the network would favor the homogenization of states due to its own structure. Specifically, we expect that the time-to-equilibrium will depend on the algebraic connectivity of the spliceosome network, *λ*_*L*_, and the number of functional proteins will decay with the *t^e^λ_L_^^*, in which *t* is time.

We tested if our two predictions, namely (*i*) the mean-field approximation of the model (equation 3) and (*ii*) the algebraic connectivity predicts the rate of loss of functional proteins across time, hold in simulations parameterized with information of the structure of empirical spliceosome network. In our simulations, at *t*=0, all proteins are functional, i.e., *ϕ*_*i*_ = *1* for any protein *i*. At the equilibrium, all states converge to zero, leading to the complete collapse of the spliceosome. We ran 1,000 simulations of the model and computed the mean number of functional proteins in a given time step and assuming *c* = *0.1* (Fig. 4b). We fit a log-log model to estimate the rate of the decay of functional proteins. To reduce statistical fluctuations we truncated the analysis for time steps in which the mean number of functional proteins is higher than one. Deviations between simulations and the analytical predictions are expected due to (1) small system size (103 proteins), (2) the difference between the binary nature of the simulation (functional vs nonfunctional proteins, equation 2) and the continuous, mean-field analytical predictions, and (3) and the loss of information on network structure due to the use of just a single eigenvalue^43^. However, the analytical predictions based on our mean-field approximation reproduced the transient dynamics of protein failures (Figure 4b). Moreover, the rate of decayment in the number of functional protein was similar to the prediction based on the algebraic connectivity (numerical simulations: *t*^*−1.80*^, mean-field prediction: *t*^*−1.86*^, algebraic connectivity: *t^e^λ_L_^^* = *t*^*−1.99*^). The rate of decay is much faster than predicted for a random network, which leads to exponential decay, *e*^*−0.54t*^, Figure 4b). Similarly, a more conservative null model that maintains the heterogeneity in the number of interactions per protein led to an exponential decay in the number of functional proteins with time (Supplementary Information). Thus, the slow decay in the number of functional proteins is not just the result of the distribution of interactions per protein. Taken together, these results suggest that the modular structure of the interactions among spliceosomal proteins lead to a network in which modules of proteins are loosely connected by interaction paths, making the system much less prone to cascading effects across the whole networks than expected for random networks.

## 6. Discussion

A network organization implies the formation of paths connecting otherwise isolated elements of the system^44^. Paths create routes for cascading effects to propagate through the system, coupling the dynamics of non-interacting elements across multiple levels of biological organization^27,45-47^. In protein networks, these paths of interactions are fundamental to a series of intracellular processes with functional consequences for the cell^48^. Previous work shows that modularity improves the robustness of networks against cascading effects^49-51^. Accordingly, spectral graph theory may provide information on the robustness of cascading effects^52,53^. Here, we integrated the approaches derived from spectral graph theory and the study of network flow to explore the consequences of the modular structure on the cascading effects in the spliceosome network. In this sense, our work improves the understanding of the vulnerability of this protein-protein network against perturbations in three main ways.

First, the network organization of the spliceosome favors the proliferation of paths. This feature leads to more paths of a given length than expected by similar-sized random networks. We show that the number of paths of a given length connecting proteins is predicted by the leading eigenvalue of the adjacency matrix describing the spliceosome. The higher proliferation rate in the spliceosome network is a consequence of the large variation in the number of interactions per protein^31^. One key finding using spectral graph analysis is that the upper bound value of the largest leading eigenvalue is the largest number of interactions recorded for a protein in a network. Biologically, the large variation in the number of interactions per protein implies that some highly connected proteins will be central for the organization of the network, creating paths among poorly connected proteins. Highly-connected proteins are central to a number of processes in the cells and deletions of such hubs in the protein networks may be lethal due to cascading effects^5^. The relevance of highly connected proteins—the centrality-lethality rule—may stem from multiple factors, such as participation in essential, direct pairwise interactions^54^ or in the organization and reorganization of the large protein network^55^. In this sense, we suggest that path proliferation is another structural consequence of highly-connected proteins that may favor cascading effects. Thus, changes in the network structure via experimental protein deletions can be predicted by analysing the spectrum (the set of eigenvalues) of the interaction matrix, which can inform the relative contribution of individual proteins to indirect paths in the protein networks. This method may improve the application of integrative approaches involving spectral graph theory and network theory to molecular biology. We know that the some of same proteins involved in the spliceosome are part of other complexes and are involved in other intracellular processes. In this sense additional insights may emerge if the structure of protein-protein interactions within the cell would be described as a multilayered network where each node (protein) can be involved in multiple subnetworks. Spectral analysis may help understanding the functional consequences of the alternative paths created by these multilayered networks^56,57^.

Second, these paths are not randomly distributed across the network—paths are, instead, concentrated within modules. Modularity is one of the main features of protein networks and evolutionary processes may favor the emergence of modularity by a combination of gene duplication, horizontal gene transfer, and natural selection^15,34^. Selection may favor modularity allowing both specificity and autonomy of functionally distinct subsets of proteins^15,35^. In this sense, the concentration of paths within modules provides a way to increase module integration, favoring distinct functional roles developed by proteins in distinct modules. In the spliceosome, modules are associated with subcomplexes that act at distinct steps of the spliceosome assembly and function^31^. Thus, path proliferation may favor the emergence of highly integrated subunits, in which effects of pairwise interactions may also activate indirect effects on non-interacting proteins associated with the same function or step of splicing process. More than promoting within-module integration, path proliferation also integrates distinct modules. There is experimental evidence that modularity does reduce the potential for cascading effects across the system^39^. However, modularity does not imply module isolation. System functioning also depends on the indirect effects between functional modules, allowing complex tasks to be completed by distinct subunits of the system. Hence, selection is not expected to favor complete module isolation, but these paths that allow system functioning may also lead to routes for cascading effects triggered by mutations and deletions. Understanding which are those paths and how they affect spliceosome functioning may be key for diagnosing diseases, and also for using specific proteins as possible pharmacological targets.

Third, the local integration of modules promoted by path proliferation and their semi-independence to other modules may provide robustness to mutations on specific proteins, as seen in human cells with SR and hnRNP families of proteins. The hnRNP proteins were previously associated with intronic miRNAs, probably facilitating splicing reactions on these pre-RNA substrates^58,59^. Some SR proteins and hnRNP-A2/B1 and hnRNP-U are modulators of splicing in SMN1 and SMN2 genes, but not other hnRNP proteins^60^. In consequence, splicing defects on SMN1 and SMN2—and the emergence of a neurodegenerative disorder such as spinal muscular atrophy—might be associated with a subset of hnRNP proteins. By constraining paths within modules, the network structure may increase the robustness of the spliceosome.

It is important to notice that the semi-independence of modules does not imply network-level robustness of the spliceosome to all types of failures and errors. Nevertheless, these results provide a theoretical benchmark that help predicting which kinds of failures are likely to cause network-level collapse in the spliceosome network. For instance, we should expect that the errors in the proteins that are simultaneously highly connected and link distinct modules will lead more often to network-level collapse of the spliceosome. For example, PRP8 is a large, highly-connected protein acting as a core component of U5 snRNP and essential for efficiency and fidelity of splicing reactions^21^. Interestingly, human PRP8 expression is reduced with an increase in cell proliferation, possibly affecting splicing globally^19^. Moreover, because the spliceosome network has a temporal structure and subunits are assembled and disassembled sequentially during the splicing reaction, we should expect that local collapse of the early subunits to join the complex is more likely to cause the largest problems with the spliceosome functioning. In fact, mutations in a group of proteins that associate during early spliceosome assembly, among which are U2AF35, SF3A1 and SF3B1, is frequently associated to development of myeloid neoplasms^61^. In this context, our study provides a network-based explanation for alterations that might lead to splicing collapse: it will be a consequence of the local collapse of modules due to cascading effects propagating across path proliferation.

The interplay between modularity and path proliferation provides an hypothesis on how protein networks preserve the interconnectivity among functional modules and constrains deleterious cascading effects to propagate across the system. Future studies should investigate the role of modular structure in robustness by combining experiments *in vivo* in which key proteins are deleted with network analysis of protein role in the organization of spliceosome network. By now, our results support that the existence of multiple, indirect paths connecting proteins is a potentially relevant consequence of the network structure for protein-protein interactions. Mapping the fragile and robust points of such networks could aid the development of new therapies, given that the misregulation of the spliceosome resulting from mutations in their proteins and from single-point mutations changing the splicing of a given gene are linked to many human diseases, such as spinal muscular atrophy, retinitis pigmentosa and several types of cancer, such as lymphocytic leukaemia and myelodysplasia^19,62^. We suggest that the approach introduced here to uncover the distribution of paths and their potential dynamical implications to spliceosome may help to characterize other types of molecular networks. In this sense, the characterization of indirect paths and their possible consequences in multiple molecular networks may provide insights on the role of the network structure in shaping the emergence of complex diseases, contributing to the emerging field of network medicine^63,64^.

## 7. Acknowledgments

We thank Ana Paula Assis, Pâmela C. Santana and Leandro Giacobelli for helpful comments. PRG was supported by CNPq and FAPESP (2017/08406-7). PPC was supported by FAPESP (2017/06994-9). MC was supported by a PMP/BS postdoctoral fellowship (UFPR/UNIVALI 46/2016).

## 8. Methods

### Spliceosome protein-protein network

The spliceosome is a macromolecular complex that is relevant to gene expression regulation^65^. Here, we briefly describe the way the spliceosome network was built up. A detailed description of sampling procedure and sensitivity analyses for different model species, data sources, interaction reliabilities is available at^31^. We used protein-protein interaction data from the spliceosome of *S. cerevisiae* available in the STRING database (http://string-db.org). The protein-protein interactions of the spliceosome can be depicted by a network encoded in an adjacency matrix **A**, in which matrix element *a_ij_* informs if protein *i* interacts with protein *j a_ij_* = *a_ji_* = *1* and zero otherwise. However, the empirical support for putative protein-protein interactions vary across pairs of proteins. In this sense, the evidence derived from multiple experiments was integrated in a score provided by the STRING database, varying from zero to one. We assumed there is a protein-protein interaction if the STRING score for the interaction was higher than a given cutoff value. Previous work shows that there are two informative cutoffs, a permissive cutoff value (0.15) and a more restrictive cutoff value (0.5)^31^. We focused the more restrictive cutoff value, but all analyses assuming the more permissive cutoff value a are available in the Supporting Information.

### Paths in connecting proteins from the same or from distinct modules

We computed 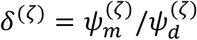, in which 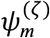 is mean number of paths with length *ζ* starting and ending with proteins from the same module and 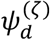 is the mean number of paths with length *ζ* starting and ending with proteins from distinct modules. We first assigned proteins to distinct modules by finding a module partition that maximizes the metric *Q ^9^* under a simulated annealing algorithm (for more details of the module structure of the spliceosome network please refer to^31^. We then computed 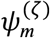 by dividing the number of paths with length *ζ* starting and ending with proteins from the same module by the number of pairs of proteins assigned to the same module. Accordingly, we computed 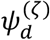 by dividing the number of paths with length *ζ* starting and ending with proteins from different modules by the number of pairs of proteins assigned to different modules.

### Simulation of cascading effects

We assigned to each protein *i* a state *ϕ*_*i*_ and we assumed *c* = *0.1*. At timestep *t*=0, we assumed that all proteins were functional, *ϕ_i_* = *1* for any protein *i*. Then, we randomly selected a protein to become non-functional, *ϕ*_*i*_ = 0. At each timestep, the probability of functional protein to become non-functional was defined by equation (2). The simulation proceed until all proteins became non-functional. We ran 1,000 simulations for the empirical network and we recorded the median number of functional proteins in each timestep. We used the same simulation algorithm to the theoretical networks generated by each of the two null models (see below) used in the manuscript (one simulation per network; 1,000 networks generated by each null model).

### Null models

We used two null models to explore the role of network structure in fueling or inhibiting cascading effects. Our first null model is the Erdos-Renyi random graph. For each pair of potentially interacting proteins we sample a random number from an uniform distribution U[0,1] and if this number was smaller than the connectance, which is the fraction of all possible interactions actually recorded in the spliceosome network, we assign an interaction. This null model generates networks with the same number of proteins and similar number of interactions to those observed in the spliceosome network, but without any nonrandom structural pattern. We also used a second null model. In this model, we also preserved the heterogeneity in the number of interactions of the spliceosome network by assuming the probability of two proteins interact is proportional to 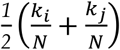, in which *k_i_*(*k_j_*) is the number of proteins interacting with protein *i* (*j*) and *N* is the number of proteins in the network. Therefore, each pair of proteins has a given probability to interact. Again, for each pair of potentially interacting proteins, we sampled a random number from an uniform distribution U[0,1] and if the sampled number was smaller than the interaction probability we assigned an interaction. We generated 1000 replicates of each null model. The MATLAB scripts used to generated the null model networks and all the analyses are available upon request.

